# Differential heating of metal nanostructures by radio frequencies: a theoretical study

**DOI:** 10.1101/2021.03.31.437926

**Authors:** Nicholas J. Rommelfanger, Zihao Ou, Carl H.C. Keck, Guosong Hong

## Abstract

Nanoparticles with strong absorption of incident radio frequency (RF) or microwave irradiation are desirable for remote hyperthermia treatments. While controversy has surrounded the absorption properties of spherical metallic nanoparticles, other geometries such as prolate and oblate spheroids have not received sufficient attention for application in hyperthermia therapies. Here, we use the electrostatic approximation to calculate the relative absorption ratio of metallic nanoparticles in various biological tissues. We consider a broad parameter space, sweeping across frequencies from 1 MHz to 10 GHz, while also tuning the nanoparticle dimensions from spheres to high-aspect-ratio spheroids approximating nanowires and nanodiscs. We find that while spherical metallic nanoparticles do not offer differential heating in tissue, large absorption cross sections can be obtained from long prolate spheroids, while thin oblate spheroids offer minor potential for absorption. Our results suggest that metallic nanowires should be considered for RF- and microwave-based wireless hyperthermia treatments in many tissues going forward.

## Introduction

Radio frequencies (RF) and microwaves are used in the clinic for emerging hyperthermia treatments to destroy cancerous growths in tissues throughout the body [1–3]. Hyperthermia can be applied as a standalone treatment [3], or it can be delivered as an adjuvant therapy in conjunction with chemotherapy [2] and/or radiotherapy [4]. While the high tissue penetration of RF/microwaves enables non-invasive therapies via antenna arrays surrounding the body [2], the achievable spatial focusing is fundamentally limited by the long wavelengths of RF. Thus, others have turned to implantable microwave antennas to minimize off-target heating [1], although this technique is more invasive.

Nanomaterials offer new capabilities to “focus” energy delivery in biological systems (Figure 1A) [5,6]. In particular, due to well-established synthesis techniques, tunable localized surface plasmon resonance (LSPR), and high biocompatibility, gold nanostructures have found widespread use in biomedical applications. Plasmon-resonant gold nanorods are highly efficient at converting near-infrared (NIR) light into heat [7] and have shown promise for treatment of shallow tumors *in vivo* [8]. Recently, programmable photothermal gene editing was achieved by second near infrared (NIR-II) illumination of a polymer-coated gold nanorod paired with a Cas9 plasmid that was triggered by a heat-inducible promoter [9]. Photothermal techniques are increasingly applied in experimental neuroscience, recently enabling through-scalp stimulation of deep brain regions in mice [10]. Additionally, the rapid heating of spherical gold nanoparticles under 532-nm illumination produced non-genetic, photocapacitive stimulation of both cultured neurons and mouse hippocampal slices [11], while gold nanorods conjugated to temperature-sensitive ion channels in the retina endowed NIR sensitivity in mice and in post-mortem human retinas [12].

**Figure 1.**
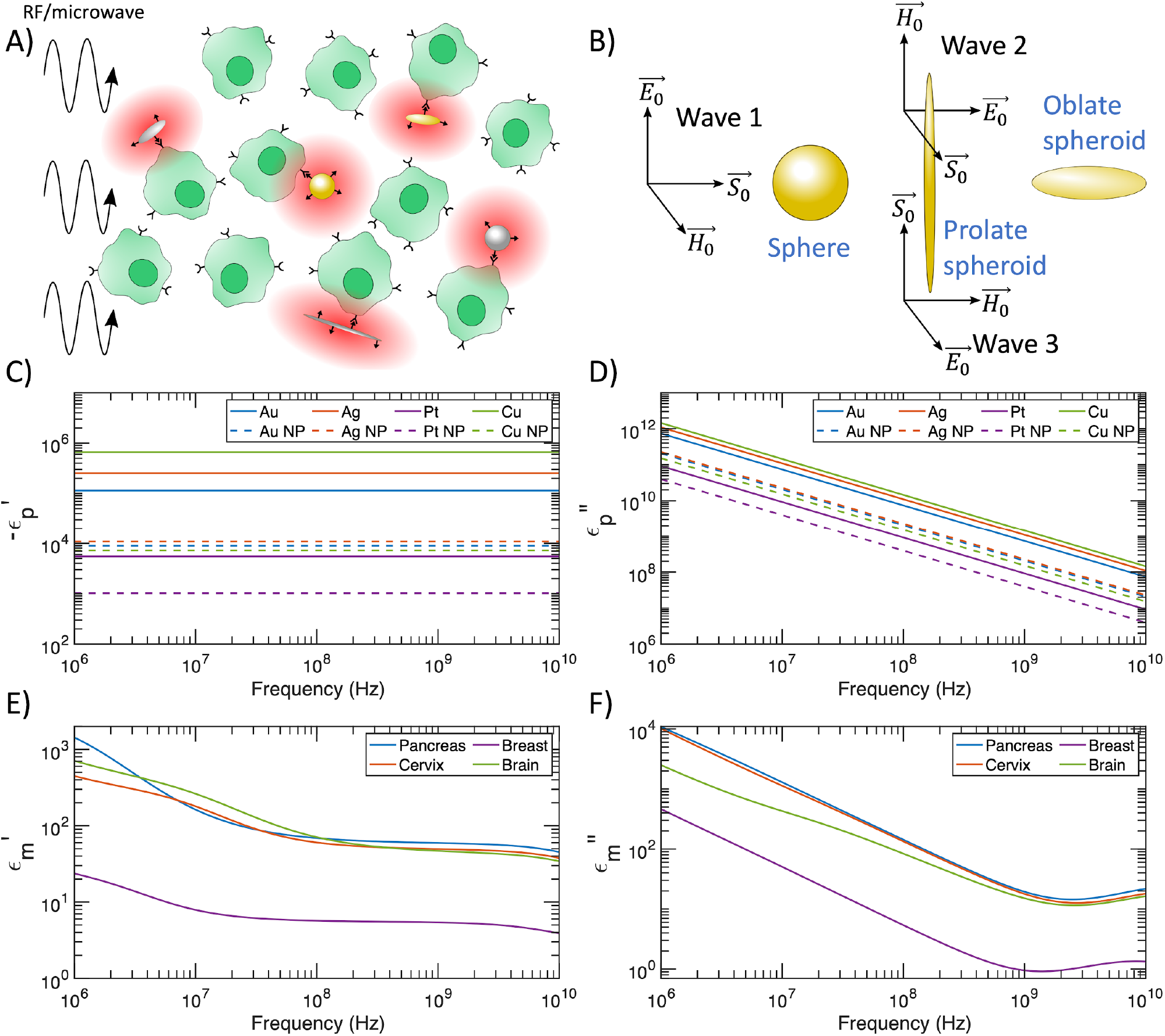
A) Schematic of differential heating induced by RF/microwave application to cells targeted by metal nanostructures. B) Orientation of RF/microwave interaction with nanomaterials. The relative absorption ratio is averaged over the 3 propagating waves shown (with equal magnitudes) to account for a collection of randomly oriented spheroids. C) Negative real component of the dielectric function for metals considered in this manuscript. Dashed curves include scattering rate corrections for spherical nanoparticles with 10 nm radius (NP = nanoparticle). D) Imaginary component of the dielectric function for metals considered in this manuscript. Dashed curves include scattering rate corrections for spherical nanoparticles with 10 nm radius. E) Real component of the dielectric function for tissues considered in this manuscript. Data from reference [37]. F) Imaginary component of the dielectric function for tissues considered in this manuscript. Data from reference [37].

Radio frequencies offer deeper tissue penetration than infrared light, but the interaction between metal nanostructures and RF/microwaves is less understood and has been the subject of controversy in recent years [13]. Initially, several studies reported heating of spherical gold nanoparticle suspensions in RF fields [14–18]. However, these reports were contradicted by subsequent experiments that discovered Joule heating of the stock solution caused the measured temperature increase of a gold nanoparticle suspension under RF irradiation [19,20]. Additionally, a theoretical investigation analyzed several classical and quantum absorption mechanisms for metallic nanoparticles and found that no combination of the considered mechanisms could produce the previously reported heating [21]. However, for nanoparticles under 10 nm in diameter, an electrophoretic mechanism was later demonstrated both experimentally [22] and theoretically [23] to account for the observed heating in gold nanoparticle suspensions. A more recent theoretical work suggested that electrophoretic resonances of small (< 5 nm) gold nanoparticles in the microwave regime would lead to promising differential heating in a weak electrolyte solution [24]. Several key reviews detail the state of the field for heating spherical gold nanoparticles with RF [25,26].

Although opportunities for rapid heating of spherical metallic nanoparticles in RF appear limited to electrophoretic heating, alternative nanoparticle geometries may offer opportunities for significant differential heating in tissue. A theoretical investigation predicted strong local field enhancements in the near-field zone around carbon nanotubes (CNTs) [27], and differential heating was indeed observed in an experimental report that exposed a tissue-mimicking phantom with CNTs to microwaves [28]. This so-called “lightning rod effect” was also predicted to occur for elongated ellipsoids [21], and a recent publication found that the differential heating of silver nanowires under RF increased with increasing aspect ratio, while confirming that spherical silver nanoparticles did not heat under identical conditions [29]. Another recent work reported that gold nanowires longer than 6 µm absorb microwave radiation [30]. A key theoretical investigation based on the electrostatic approximation [31] derived the absorption cross section for an ellipsoidal particle, followed by numerical examples considering an electrophoretic heating mechanism valid for small nanoparticles [32]. While Joule heating of small spherical gold nanoparticles in RF has been previously discounted [21], the authors of this ellipsoidal investigation did not consider the potential for meaningful Joule heating in larger ellipsoidal particles. The high-aspect-ratio limit of a prolate spheroid is a nanowire (Figure 1B), which are easily synthesized in the laboratory and have recently shown promise for differential heating [29]. Likewise, the high-aspect-ratio limit of an oblate spheroid is a nanodisc (Figure 1B), which may also demonstrate promising absorption characteristics. Further, while most studies have investigated spherical nanoparticles on the 1 nm - 100 nm lengthscale, larger particles in the 100 nm - 10 µm lengthscale are still injectable and could be used safely *in vivo* for hyperthermia treatments [33]. Lastly, a study that compares the differential heating of different types of metal nanoparticles in various tissues is missing from the literature, and such a study is important for optimizing hyperthermia treatment protocols.

In this manuscript, we present calculations for spherical and spheroidal metal nanoparticles in a variety of tissues that have been commonly treated with RF and/or microwave hyperthermia in clinical trials [1,2]. We consider particles ranging in size from nanometers to tens of microns across the frequency spectrum from 1 MHz to 10 GHz, covering the operating frequencies of nearly all hyperthermia treatments [1,2]. The main novelty of this work over existing studies is three-fold: first, when considering the electronic response of nanoparticles to incident radiation, we scan over a parameter space of realistic values that can be achieved by synthesizing metal nanostructures with reported methods. Second, we use different types of biological tissue with reported complex dielectric constants for calculating the differential heating of nanoparticles placed therein, instead of an aqueous electrolyte solution as the model for the lossy media. Third, our paper, for the first time, proves that metal nanostructures of realistic electrical and geometrical properties can achieve significant differential heating over biological tissues in RF and microwave fields. We hope this theoretical investigation helps to explain previous experimental results, while also suggesting promising particle geometries and frequencies to optimize future hyperthermia protocols.

### Theory

After establishing the notational conventions of this manuscript and the methods used to obtain dielectric functions for metallic nanoparticles and biological tissues, we consider the power absorbed by an ellipsoidal particle under the electrostatic approximation, following references [31,32]. We observe that in the special case of a spherical particle, the result matches the absorption cross obtained from Mie theory. We conclude with a discussion on the regime of validity of the electrostatic approximation.

## I. Conventions

Let *μ*_0_ and *ϵ*_0_ denote the permeability and permittivity of vacuum, *ϵ*_*p*_ denote the complex relative permittivity of a metallic particle, and *ϵ*_*m*_ denote the complex relative permittivity of the surrounding medium. We follow the convention *e*^−*iωt*^ for time harmonic fields; thus, the imaginary part of the complex relative permittivity for a passive material is positive: *ϵ* = *ϵ*′ + *iϵ*″. The wavenumber of vacuum is given by 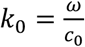, where *c*_0_is the speed of light in vacuum, the complex wavenumber of a metallic particle is given by 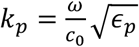, and the complex wavenumber of the surrounding medium is given by 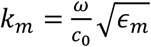. We define the complex index of refraction as 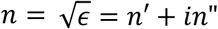. Thus, we have 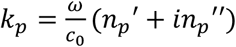and 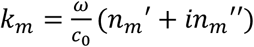.

## II. Dielectric functions of metal nanoparticles and tissue

We calculate the dielectric function of metal nanoparticles using the Drude model for free electrons:

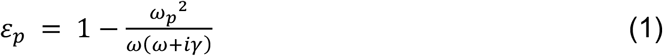

where *ω*_*p*_ is the plasma frequency and *γ* is the scattering rate [31]. The Drude model is a good fit to the dielectric function of metals at low frequencies [34], appropriately producing the DC conductivity as *ω* → 0 (please see **Supplementary Information** for derivation). The plasma frequency *ω*_*p*_ is defined as:

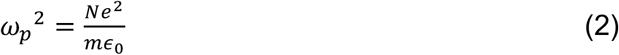

where *N* is the number density of free electrons, *e* is the charge of an electron, and *m* is the effective mass of an electron [31].

The scattering rate *γ* is dependent on electron scattering from impurities, lattice defects, other electrons, and phonons [35]. In the classical treatment, the scattering rate can be calculated as *γ*_*bulk*_ = *v*_*F*_/*L*_∞_, where *v*_*F*_ is the Fermi velocity and *L*_∞_ is the mean free path in the bulk metal [35]. However, in nanoparticles smaller than the electron mean free path, changes to the scattering rate are required to match experimental measurements of small nanoparticles [36]. The formula for *γ* can be modified by introducing a size-dependent term to account for electron scattering from the nanoparticle surface [35,36]:

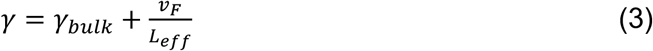

where *L*_*eff*_ is the effective mean free path. Although there is some disagreement, *L*_*eff*_ = 4*R*/3 is typically used for spherical nanoparticles of radius *R* [31]. An in-depth analysis relying on geometrical probability calculated *L*_*eff*_ for spheres, prolate spheroids, and oblate spheroids [35], and we use these relations when calculating the scattering rate *γ* in this manuscript. For a sphere of radius *R*, the effective mean free path is given by [35]:

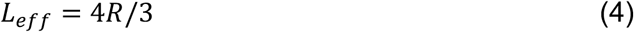

matching the typical value reported above. Next, for a prolate spheroid with major axis *L* (length) and minor axis *D* (diameter), the effective mean free path is given by [35]:

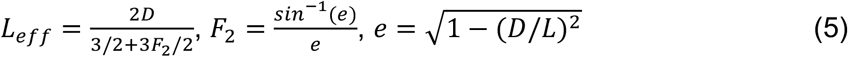

We see that as we approach the limit of a sphere (*D*/*L* → 1), *F*_2_ → 1 and thus *L*_*eff*_ → 4*R*/3 as expected. Lastly, for an oblate spheroid with major axis *D* (diameter) and minor axis *T* (thickness), the effective mean free path is given by [35]:

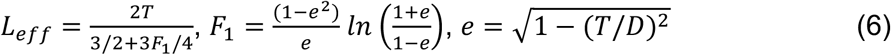

Once again, in the limit of a spherical particle (*T*/*D* → 1), *F*_1_ → 2 and thus *L*_*eff*_ → 4*R*/3 as expected.

We apply scattering rate corrections for all particle sizes considered in our analysis, rather than cutting off the correction for particles with *L*_*eff*_ > *L*_∞_, as it is reasonable to assume that particles slightly larger than the bulk mean free path will still host increased scattering rates due to the finite particle size. The correction becomes insignificant for particles with *L*_eff_ ≫ *L*_∞_, as we would expect from the underlying physics.

We extract the dielectric function *ϵ*_*m*_ for different biological tissues across the 1 MHz - 10 GHz frequency range from the Gabriel parameterizations [37–42]. In particular, we consider the Gabriel parameterizations for pancreatic, cervical, breast, and brain tissue, as these tissues are frequent targets of RF or microwave hyperthermia treatments in clinical trials [1,2]. For brain tissue, we use a grey matter to white matter ratio of 3:2 [43]. We also consider breast and brain cancerous tissues, with dielectric properties reported in references [44,45]. Although the dielectric properties of some tissues at the cellular scale are heterogeneous and anisotropic [46], a specific challenge of performing simulation at the cellular scale arises from the limited availability of dielectric measurements of the plasma membrane at the frequency range of interest. Therefore, we apply bulk tissue dielectric measurements throughout this manuscript to increase the tractability of our simulations and the generalizability of our results.

## III. Absorption cross section of an ellipsoid

We will consider an ellipsoidal particle placed within a uniform electrostatic field 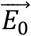 the following section for constraints on the validity of this approximation). Reference [31] solved the problem of an ellipsoidal particle placed in an electrostatic field 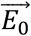 applied parallel to the *z*- axis (length of *x* semi-axis of ellipsoid = *a*, length of *y* semi-axis of ellipsoid = *b*, length of *z* semi-axis of ellipsoid = *c*), finding that the electric field 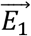 produced within the particle is both constant and parallel to 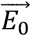. The authors found that the asymptotic perturbing potential induced by the presence of the ellipsoidal particle matches the potential of a dipole with dipole moment also in the *z*-axis:

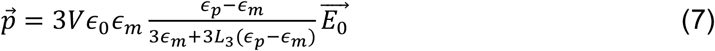

where V is the volume of the ellipsoid given by 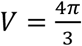, and *L*_3_ is given by:

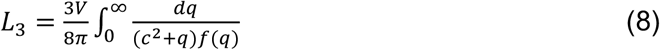

where 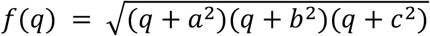. On the other hand, the polarizability *α*_*j*_ of a particle in any of the three Cartesian axes is defined by [31]:

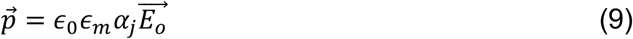

Here, we notice that we can extract the polarizability *α*_3_, which is that in the *z*-direction, by comparing Equation (7) and Equation (9). In doing so, we can also generalize our results to an electrostatic field applied along any of the 3 principal axes, as the *z*-axis we initially chose holds no special properties. We fix the semi-axes to be parallel to the cartesian axes, given by 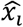 for *i* =1, 2, and 3. Thus, for an applied electrostatic field 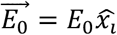with *i* =1, 2, or 3 we have:

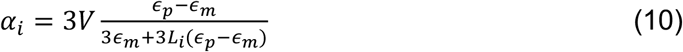

with

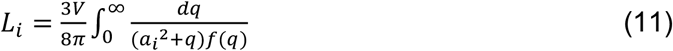

where *a*_1_ = *a, a*_2_ = *b*, and *a*_3_ = *c* per the definitions given at the beginning of this section. The geometrical factors *L*_1_, *L*_2_, and *L*_3_ sum to 1 for all ellipsoidal geometries. Next, following the derivation in reference [32], we see that the field within the particle 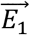 is given by:

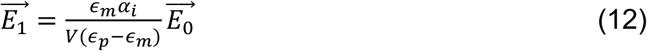

Assuming the particle’s dielectric constant is uniform throughout its volume, we can solve for the power loss within the particle using Poynting’s theorem:

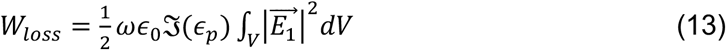

From Equation (12), we see that the electric field within the particle is uniform, simplifying the volume integration:

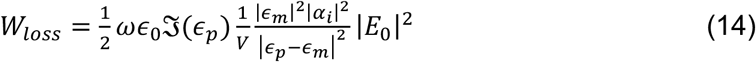

We can arrive at the absorption cross section *C*_*abs*_ by normalizing *W*_*loss*_ by the power density *P* of the incident plane wave:

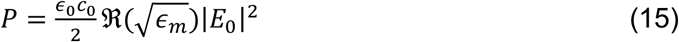

and thus:

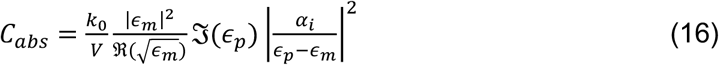

Substitution of *α*_*i*_ and subsequent simplification gives:

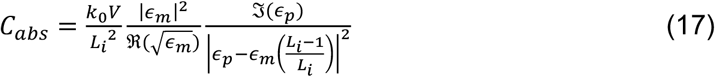

Here we note that for the special case of a sphere, 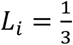 for *i* =1, 2, and 3. Substituting 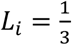 into Equation (17) for a sphere of radius *r*, we reach:

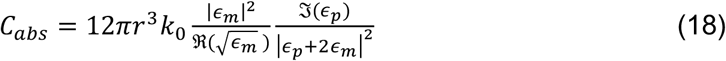

which matches the absorption cross section for a sphere in an absorbing medium derived in reference [23] with an independent approach based on Mie theory.

## IV. Validity bounds of the electrostatic approximation

The electrostatic approximation offers a useful simplification to calculate absorption cross sections over the broad parameter space covered in the plots below. Rather than relying on computationally expensive numerical techniques, the analytic expressions presented above can easily be evaluated by users in the laboratory or clinic to gain an approximate understanding of the differential heating of a particular combination of frequency, tissue type, and nanoparticle composition, shape, and size. In this section, we consider the physical validity of the electrostatic approximation, and review appropriate bounds to govern its application with RF/microwave fields.

Consider an ellipsoidal particle illuminated by a monochromatic plane wave of the form 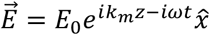. At a particular moment in time, the electric field amplitude illuminating the spheroid is given by 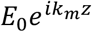. If the longest semi-axis of the spheroid (with length *a*) is oriented along the *z*-direction, then for 𝔍 (*k*_*m*_*a*) ≪ 1 and ℜ (*k*_*m*_*a*) ≪ 1, we have 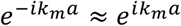, and thus the particle is approximately exposed to a uniform field across the volume it occupies. This sets the first condition for validity of the electrostatic approximation:

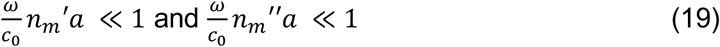

Bohren and Huffman showed that the field within an ellipsoidal particle exposed to a uniform electrostatic field aligned with one of its axes is uniform and parallel to the incident field [31]. However, to ensure that the field within the particle is also uniform when exposed to a *plane wave*, we require:

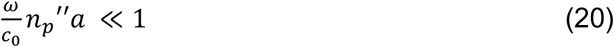

Lastly, considering that the field of the incident plane wave changes with a characteristic time of order *τ* = 1/*ω* and that it takes approximately *τ*^*^ = *an*_*p*_′/*c*_*o*_ for signal propagation across the ellipsoid, we desire *τ*^*^ ≪ *τ* in order for all points within the ellipsoid to simultaneously receive the signal of the changing field. Thus, the final condition for the electrostatic approximation is given by:

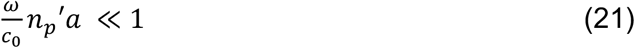

If all of these conditions are met for a given frequency, particle composition, particle size, and medium composition, then the approximation of an electrostatic field that we made in the previous section is valid. In the plots that follow, we define 0.1 as the maximum value satisfying ≪ 1.

We note that the use of the electrostatic approximation does not take into account structural features such as surface or volume inhomogeneities which could alter the absorption properties of affected nanoparticles. Specifically, polycrystalline metals are less conductive than their single-crystal counterparts due to grain boundaries, thereby leading to a higher scattering rate *γ*, a lower conductivity *σ*, and thus a lower *ϵ*_*p*_″ [47]. According to Equation (18), a higher absorption cross section *C*_*abs*_ and a higher degree of differential heating are expected as a result of polycrystallinity, grain boundaries, and other defects in spherical nanoparticles. The effects of nanoparticle coatings are also beyond the scope of the electrostatic approximation and will not be considered in this manuscript.

We also note that our analysis relying on the electrostatic approximation strictly considers isolated particles, and that the absorption properties of nanoparticle aggregates must be treated with different methods. For example, a self-assembled chain of small Au spheres may be approximated with a single prolate spheroid if the interparticle distance and chemistry affords efficient charge transfer [48–50]. However, more complex numerical simulations are needed for solving the differential heating of these self-assembled nanostructures to account for the different dielectric functions of surface-coating molecules from the conductive particle materials. Specifically, surface-coating molecules have been reported as a dielectric layer with low conductivity, thus breaking a single continuous, conductive structure into separate dipoles under external fields [51]. This discontinuity in the conductive chain cannot be properly modeled with our analytical method, thereby necessitating numerical simulations.

## V. Relative absorption ratio for randomly oriented particles

In this manuscript, we seek to provide appropriate results for *in vivo* conditions. Assuming that no alignment mechanism is present, a large collection of injected or intravenously delivered particles will be randomly oriented. Thus, our quantity of interest is the average absorption cross section < *C*_*abs*_ >, averaged over all possible particle orientations. Because we have chosen to work with plane monochromatic waves, we can express an arbitrary field as the superposition of field components that are aligned with our semi-axes. Bohren and Huffman found that the average absorption cross section of an ellipsoid is given by the average of the three absorption cross sections calculated by iteratively assuming that the incident field is parallel to the three particle semi-axes *i* for *i* =1, 2, and 3 [31]:

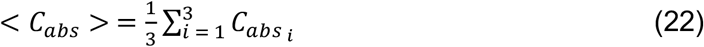

While following this approach, we calculate whether the electrostatic approximation is satisfied for each of the three wave directions shown in Figure 1B. If a particular semi-axis of a particle violates the electrostatic approximation, we set 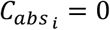 for that semi-axis *i*. Thus, < *C*_*abs*_ > calculated in this manuscript is a lower bound on the average absorption cross section, as nanostructures that violate the electrostatic approximation in particular dimensions will still absorb part of the incident radiation, in reality. We justify neglecting more complicated calculations beyond the electrostatic approximation by arguing that field strength decay within large nanoparticles is undesirable, as regions of the nanoparticle exposed to weaker fields will merely act as heat sinks and reduce the particle’s differential heating potential. The bounds of the electrostatic approximation above limit the allowable field decay within nanoparticles, so by remaining within the bounds of the electrostatic approximation, we can ensure that the entire volume of the nanoparticles under investigation will meaningfully contribute to differential heating.

We use the relative absorption ratio *F*_*abs*_ as a measure of differential heating, defined here as:

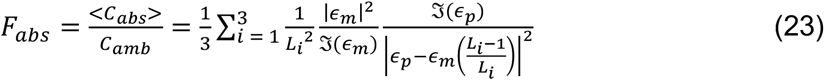

where 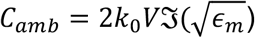 is the absorption cross section of an identical volume *V* of the tissue itself. Specifically, differential heating will occur for *F*_*abs*_ > 1. By maximizing *F*_*abs*_, the RF/microwave power necessary to achieve hyperthermic temperatures can be minimized, reducing off-target heating of healthy tissues.

## Results

The negative real part and the imaginary part of the dielectric function for the metals considered in this manuscript are plotted in Figures 1C and 1D, respectively. We use the Drude model for free electrons to calculate these curves, relying on the input parameters *ω*_*p*_ and *γ*. In the **Supplementary Information**, we discuss variation in *ω*_*p*_ and *γ* in the reported literature, and we find that the reported variation does not significantly affect our calculated relative absorption ratios. The dielectric functions for spherical nanoparticles with *R* = 10 nm are also displayed in Figures 1C and 1D, which include a correction to the scattering rate based on the effective mean free path (Equation 3). The finite size of these nanoparticles increases the scattering rate, which serves to decrease the magnitude of both *ϵ*_*p*_′ and *ϵ*_*p*_″ at the low frequencies considered in this manuscript (please see **Supplementary Information** for low frequency derivation).

Figures 1E and 1F show the real and imaginary parts of the dielectric function for the tissues considered in this manuscript, calculated based on parameterizations of Gabriel’s measurements [37–40]. While the dielectric functions for pancreatic, cervical, and brain tissue are comparable across the 1 MHz - 10 GHz spectrum, the dielectric function of the breast is significantly different due to the high lipid content of breast tissue [52,53].

### Spherical metal nanoparticles do not produce differential heating in RF fields

We begin our analysis by calculating the relative absorption ratio *F*_*abs*_ for spherical metallic nanoparticles in RF/microwave fields, a subject of controversy in recent years [13,18,19,21,23]. For the case of a spherical particle, the absorption cross section of an ellipsoid reduces to Equation (18). In Figure 2A, we list parameters that can be tuned when calculating *F*_*abs*_, namely the frequency (*f*) of the incident electromagnetic field, the radius (*R*) of the spherical nanoparticle, the type of metal (determines *ϵ*_*p*_), and the type of background tissue (determines *ϵ*_*m*_). We begin by plotting *F*_*abs*_ for Au, Ag, Pt, and Cu nanoparticles in the pancreas in Figures 2B-2E, a common target for hyperthermia therapies in clinical trials. Au, Ag, Pt, and Cu nanoparticles are chosen in this study for their ease of synthesis and surface functionalization, chemical inertness, and many demonstrated applications in biomedicine [54]. By studying Equation (23) for *F*_*abs*_, we find that in the case of a sphere (*L*_*j*_ = 1/3) assuming |*ϵ*_*p*_′| ≫ |*ϵ*_*m*_′| and |*ϵ*_*p*_″| ≫ |*ϵ*_*m*_″| (valid for all tissues and metals considered here above 10 MHz):

**Figure 2.**
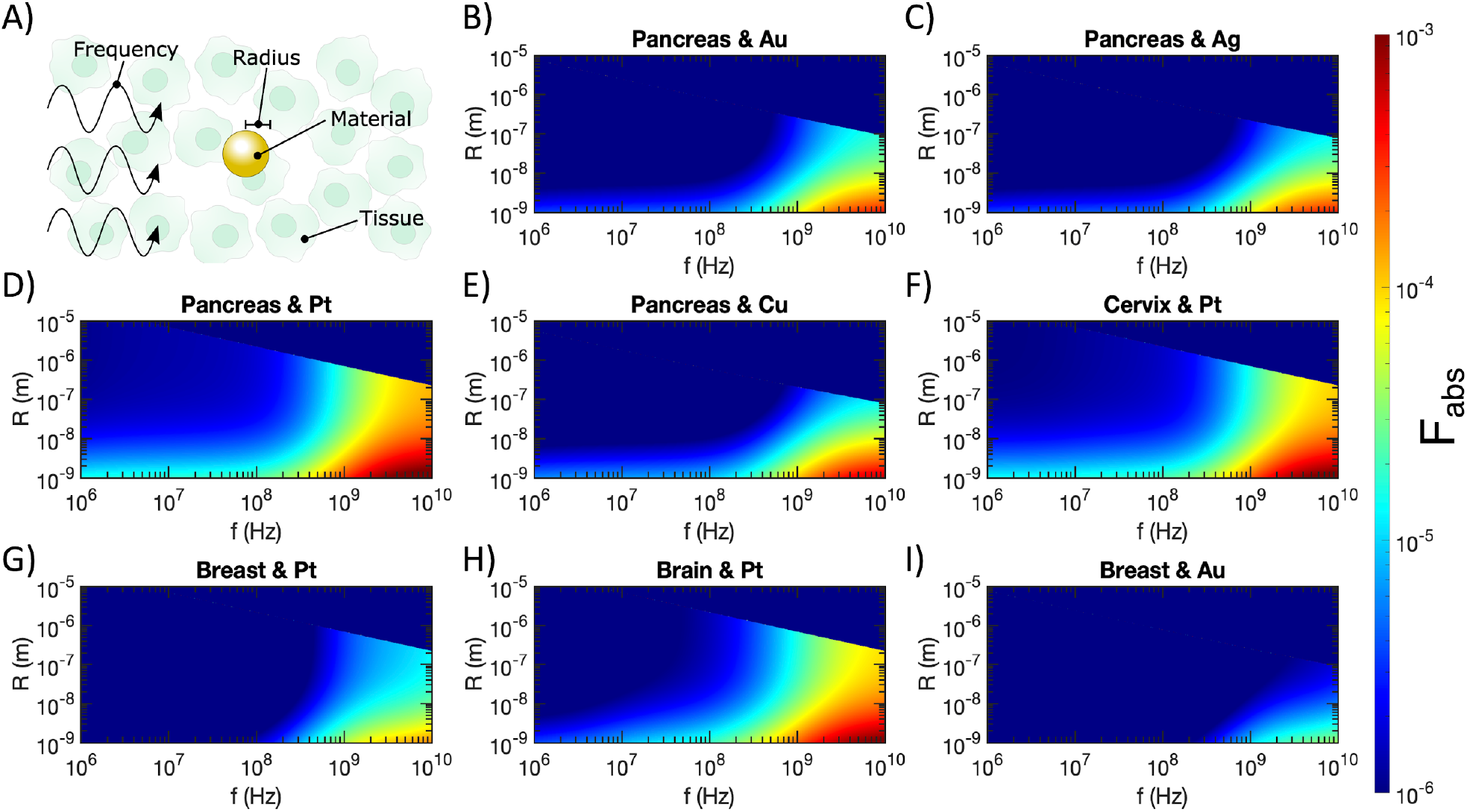
A) Labelling the tunable variables in relative absorption ratio calculations for spherical nanoparticles. B-I) Relative absorption ratio *F*_*abs*_ for metallic nanospheres in tissue (metal and tissue types marked in titles), as a function of radius (*R*) and incident electromagnetic wave frequency (*f*). Semi-axes (i.e., radii in the case of spheres) that violate the electrostatic approximation are excluded from all plots, thus these results represent lower bounds of differential heating.

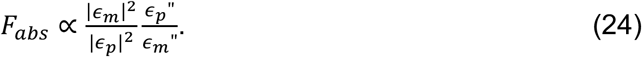

Thus, for a fixed nanoparticle material, radius, and frequency, the subsequent differential heating is determined by the ratio of |*ϵ*_*m*_|^2^/*ϵ*_*m*_″. The numerator of this ratio dominates due to the squared term, so tissues with smaller |*ϵ*_*m*_| will produce weaker differential heating, explaining the smaller *F*_*abs*_ for Pt nanoparticles in breast tissue (Figure 2G) compared to Pt nanoparticles in the pancreas, cervix, and brain tissue (Figures 2D, 2F, and 2H). Similarly, Au nanoparticles in breast tissue (Figure 2I) produce less differential heating than Au nanoparticles in the pancreas (Figure 2B).

On the other hand, for a fixed tissue type, particle radius, and frequency, the subsequent differential heating relies on the ratio *ϵ*_*p*_″/|*ϵ*_*p*_|^2^. Because the denominator dominates in this case, materials with smaller |*ϵ*_*p*_| will produce larger *F*_*abs*_. Figures 1C and 1D show that bulk Pt has the smallest |*ϵ*_*p*_| of the four metals considered, suggesting that it will produce the largest *F*_*abs*_. Indeed, we observe that Pt nanoparticles in pancreatic tissue (Figure 2D) produce a larger *F*_*abs*_ than Au (Figure 2B), Ag (Figure 2C), or Cu (Figure 2E) nanoparticles. Further, the decrease in |*ϵ*_*p*_| caused by the increase in *γ* for small nanoparticles induces a larger *F*_*abs*_, as seen throughout Figure 2. That said, even for the optimal combination of small Pt nanoparticles in breast tissue (the lowest considered |*ϵ*_*p*_| and highest considered |*ϵ*_*m*_|), *F*_*abs*_ has a maximum value of only ∼10^−3^. This aligns with previous theoretical work calculating that Joule heating is insignificant in spherical metal nanoparticles [21].

As discussed in the previous section, we do not include contributions to < *C*_*abs*_ > from ellipsoid axes that violate the electrostatic approximation. Because all three axes have identical dimensions in the case of spherical nanoparticles, *F*_*abs*_ abruptly drops to zero above a certain radius determined by *f* and *ϵ*_*p*_. Skin depth effects in these larger nanoparticles would prevent significant heating of the centers of the spheres, effectively producing a heat sink that would limit heating of surrounding tissues.

### Prolate metal spheroids produce significant differential heating in RF fields

We continue by considering prolate spheroids, which have similar geometries to nanowires in the high-aspect-ratio limit. As for the case of spherical nanoparticles, the tissue type, metal type, and frequency of the incident electromagnetic wave can be tuned, but for prolate spheroids both the diameter (*D*) and length (*L*) can be tuned (Figure 3A), adding a geometric degree of freedom compared to spheres. While *C*_*abs*_ was identical for all three axes for spherical nanoparticles, we now must consider *C*_*abs*_ contributed by unique axes for prolate spheroids and average over these cross sections to account for a collection of randomly oriented spheroids according to Equation (22). Compared to spheres, in the case of prolate or oblate spheroids, it is more challenging to predict promising metals and tissues to maximize *F*_*abs*_ based on *ϵ*_*p*_ and *ϵ*_*m*_ because *F*_*abs*_ is highly dependent on the values of *L*_G_. This parameter varies widely over the geometrical space we consider in Figures 3 and 4.

**Figure 3.**
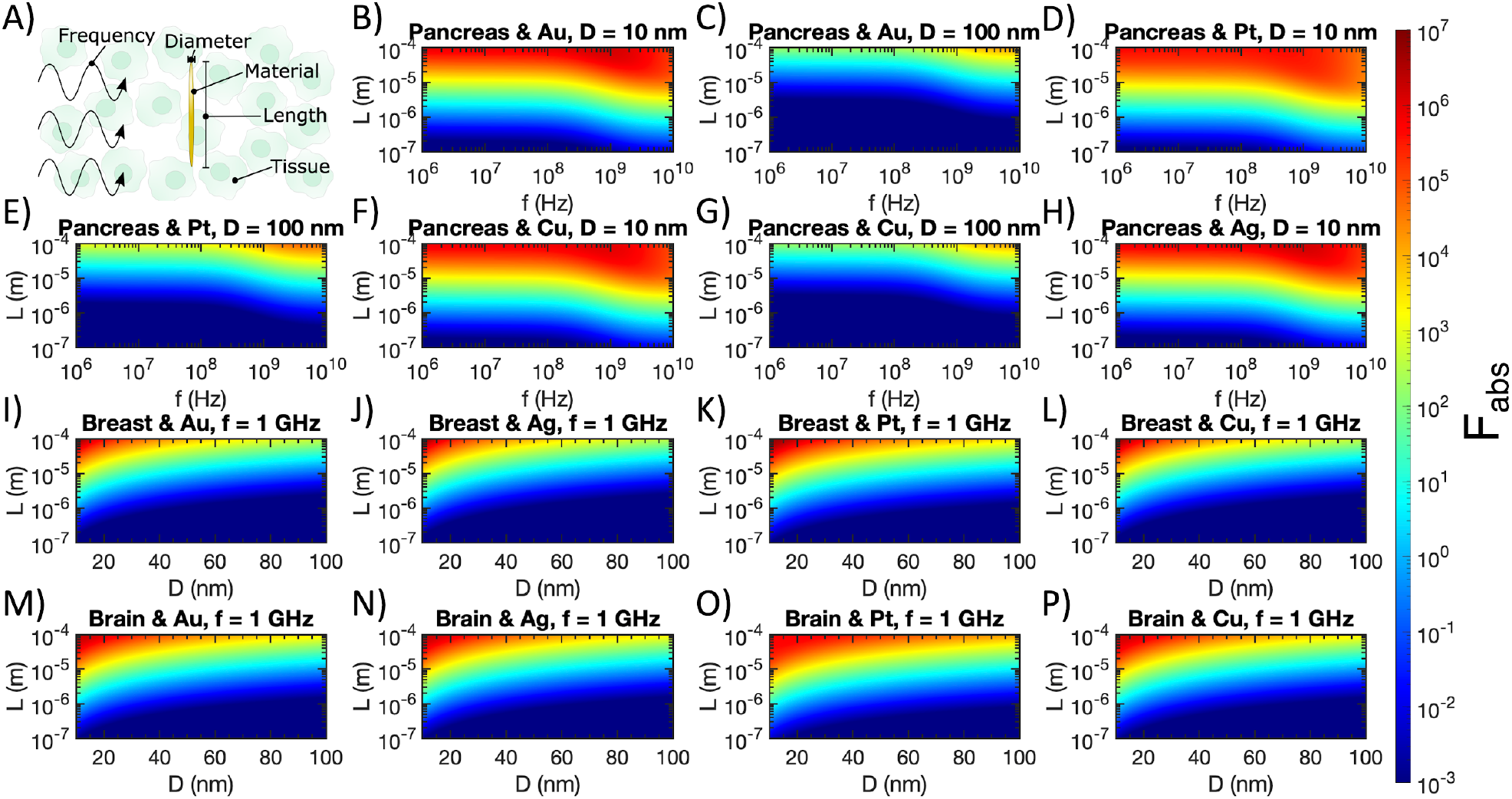
A) Labelling the tunable variables in relative absorption ratio calculations for prolate spheroids. B-H) Relative absorption ratio *F*_*abs*_ for metallic prolate spheroids in tissue (metal and tissue types marked in titles), as a function of spheroid diameter (*D*), spheroid length (*L*), and incident electromagnetic wave frequency (*f*). I-P) Relative absorption ratio *F*_*abs*_ for metallic prolate spheroids in tissue (metal and tissue types marked in titles) at *f* = 1 GHz, as a function of spheroid diameter (*D*) and spheroid length (*L*). Semi-axes that violate the electrostatic approximation are excluded from all plots, thus these results represent lower bounds of differential heating.

**Figure 4.**
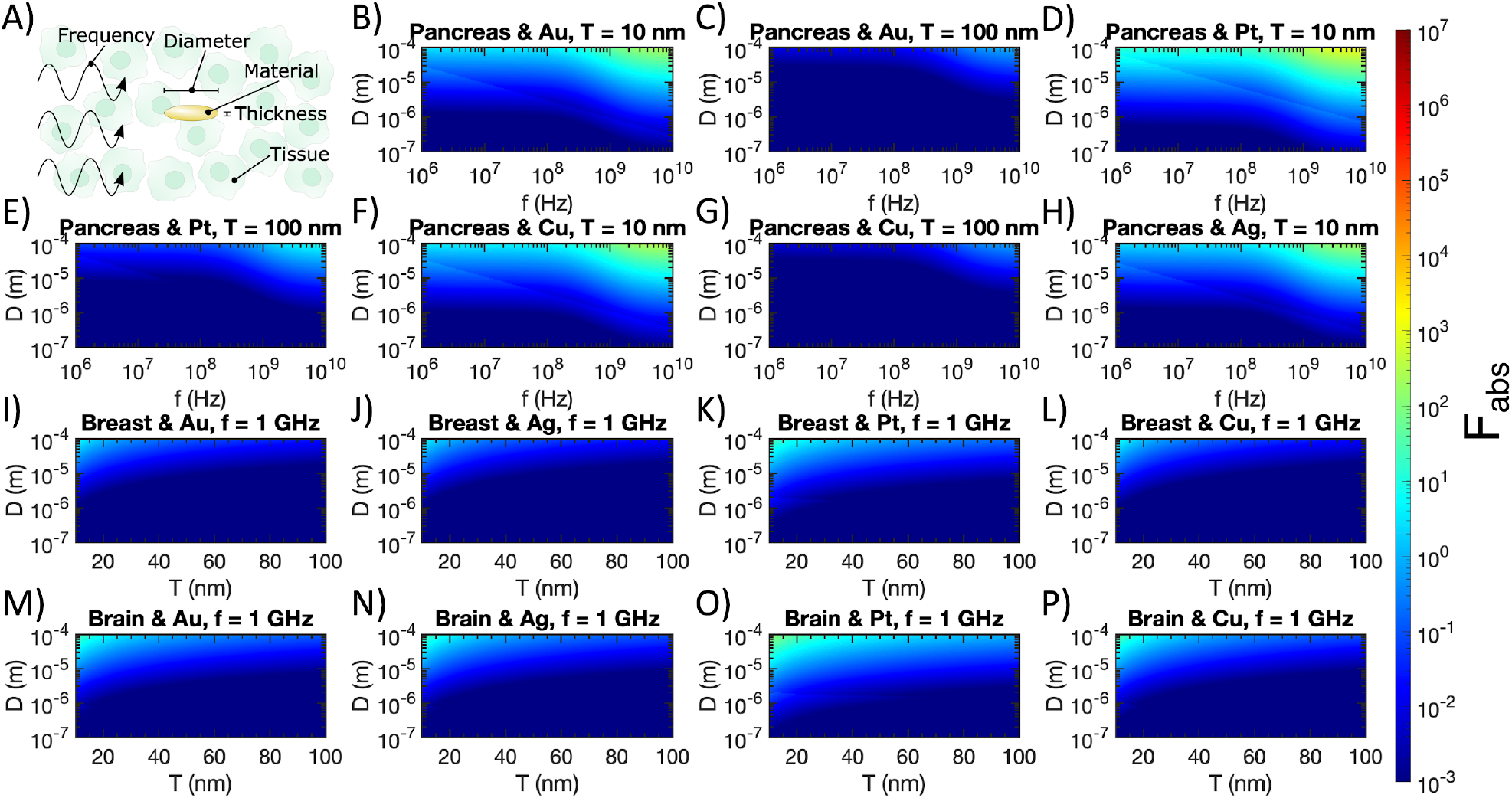
A) Labelling the tunable variables in relative absorption ratio calculations for oblate spheroids. B-H) Relative absorption ratio *F*_*abs*_ for metallic oblate spheroids in tissue (metal and tissue types marked in titles), as a function of spheroid diameter (*D*), spheroid thickness (*T*), and incident electromagnetic wave frequency (*f*). I-P) Relative absorption ratio *F*_*abs*_ for metallic oblate spheroids in tissue (metal and tissue types marked in titles) at *f* = 1 GHz, as a function of spheroid diameter (*D*) and spheroid thickness (*T*). Semi-axes that violate the electrostatic approximation are excluded from all plots, thus these results represent lower bounds of differential heating.

The upper regions of Figures 3B, 3D, 3F, and 3H consider nanowires in the high-aspect-ratio limit in the pancreas, a common target for hyperthermia therapies in clinical trials. Impressively, *F*_*abs*_ up to ∼10^7^ are attainable for prolate spheroids made of Au or Ag with *D* = 10 nm and *L* = 100 µm in the pancreas. For the prolate spheroids with *D* = 100 nm considered in Figures 3C, 3E, and 3G, we observe smaller values of *F*_*abs*_ for the same metals, frequencies, and tissue types. These spheroids have lower aspect ratios than the *D* = 10 nm case, drawing closer to the case of spherical nanoparticles which showed minimal *F*_*abs*_.

The preferable differential heating of high-aspect-ratio prolate spheroids is emphasized in Figures 3I - 3P, where it is clear that spheroids with smaller *D* and larger *L* produce the largest *F*_*abs*_ at 1 GHz. Indeed, the largest values of *F*_*abs*_ are obtained in the upper left corner (smallest *D* and largest *L*) for all tissue/metal combinations considered. In this regime, for Pt and Cu spheroids, slightly greater differential heating is obtained in breast tissue compared to brain tissue, while the opposite was observed for spherical Pt nanoparticles in Figure 2.

Violations of the electrostatic approximation are less obvious in Figure 3 than in Figure 2. This is because the largest *C*_*abs*_ is obtained when the electric field is aligned with the long axis of the prolate spheroid 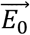 in Wave 1 in Figure 1B). Since the electric field’s amplitude oscillates over the direction of 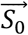 in Wave 1, the magnitude of the electric field varies minimally over the (small) diameter of the prolate spheroid. Thus, the axis with the largest contribution to < *C*_*abs*_ > is unlikely to violate the electrostatic approximation, at least for the range of diameters considered in this manuscript.

### Oblate metal spheroids produce moderate differential heating in RF fields

We conclude our analysis by considering the case of oblate spheroids, which we use to approximate nanodiscs in the high-aspect-ratio limit. As shown in Figure 4A, the diameter (*D*), thickness (*T*), and metal type of the oblate spheroids can be tuned, in addition to the tissue type and incident frequency (*f*). For the plots in Figure 4, we retain the color bar from Figure 3 to facilitate comparison between prolate and oblate spheroids. Similarly to the case of prolate spheroids, we observe larger *F*_*abs*_ for higher-aspect-ratio spheroids, as illustrated by the preferable absorption in Figures 4B, 4D, 4F, and 4H with 10 nm thickness compared to Figures 4C, 4E, and 4G with 100 nm thickness. While differential heating in the pancreas is still attainable with oblate metallic spheroids, the maximum value of *F*_*abs*_ for oblate spheroids is ∼5 orders of magnitude smaller than the case of prolate spheroids.

In Figures 4I-4P, we confirm that at 1 GHz in breast or brain tissue, thin oblate spheroids with large diameters are preferable to maximize absorption. Pt spheroids show the largest *F*_*abs*_ in both breast and brain tissue, while spheroids of all metals are more absorbing in brain tissue than breast tissue. That said, oblate spheroids perform much worse than prolate spheroids of identical aspect ratio for all metals considered here. Considering the electrostatic approximation, the largest absorption cross sections are contributed from electric fields aligned with the long axes of oblate spheroids (Wave 2 and Wave 3) in Figure 1B. While Wave 2 is likely to violate the electrostatic approximation because the amplitude of 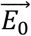 in Wave 2 varies over the long axis of the spheroid, Wave 3 is unlikely to violate the electrostatic approximation because the amplitude of 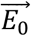 in Wave 3 varies over the (small) thickness of the spheroid. Thus, a factor of 2 in *F*_*abs*_ is lost when the electrostatic approximation is violated by Wave 2, accounting for the discrete changes in *F*_*abs*_ visible in Figure 4. Although Wave 1 is also likely to violate the electrostatic approximation, its contribution to < *C*_*abs*_ > is less significant as the electric field is aligned with the short axis of the oblate spheroid.

### Changes in absorption in cancerous tissues

While the results above were calculated using the dielectric functions of normal tissues, the dielectric functions of some cancerous tissues have been reported with larger |*ϵ*| than their healthy counterparts [44]. Thus, it is useful to calculate *F*_*abs*_ in cancerous tissues with reported dielectric functions to check how the relative absorption ratio changes. It is clear from Equation (24) that *F*_*abs*_ will increase for spherical nanoparticles in cancerous tissues with larger |*ϵ*|. The relationships for prolate and oblate spheroids are less straightforward and depend on the aspect ratio of the nanostructures. In Figure 5, we calculate *F*_*abs*_ for spherical nanoparticles (5A-5D), prolate spheroids (5E-5H), and oblate spheroids (Figure 5I-5L) at 1 GHz in breast cancer and brain cancer tissues. While we indeed calculate an increase in *F*_*abs*_ for spherical nanoparticles, this increase is still not sufficient to achieve differential heating in breast or brain cancer. The trends for prolate and oblate spheroids hold as before, with high-aspect-ratio prolate spheroids offering significant potential for differential heating (maximized by Au in brain cancer with *F*_*abs*_ ≈ 1.3 × 10^6^, slightly lower than *F*_*abs*_ ≈ 1.9 × 10^6^ for Au prolate spheroids in healthy brain tissue), while oblate spheroids offer only minor potential for differential heating in the high-aspect-ratio limit.

**Figure 5.**
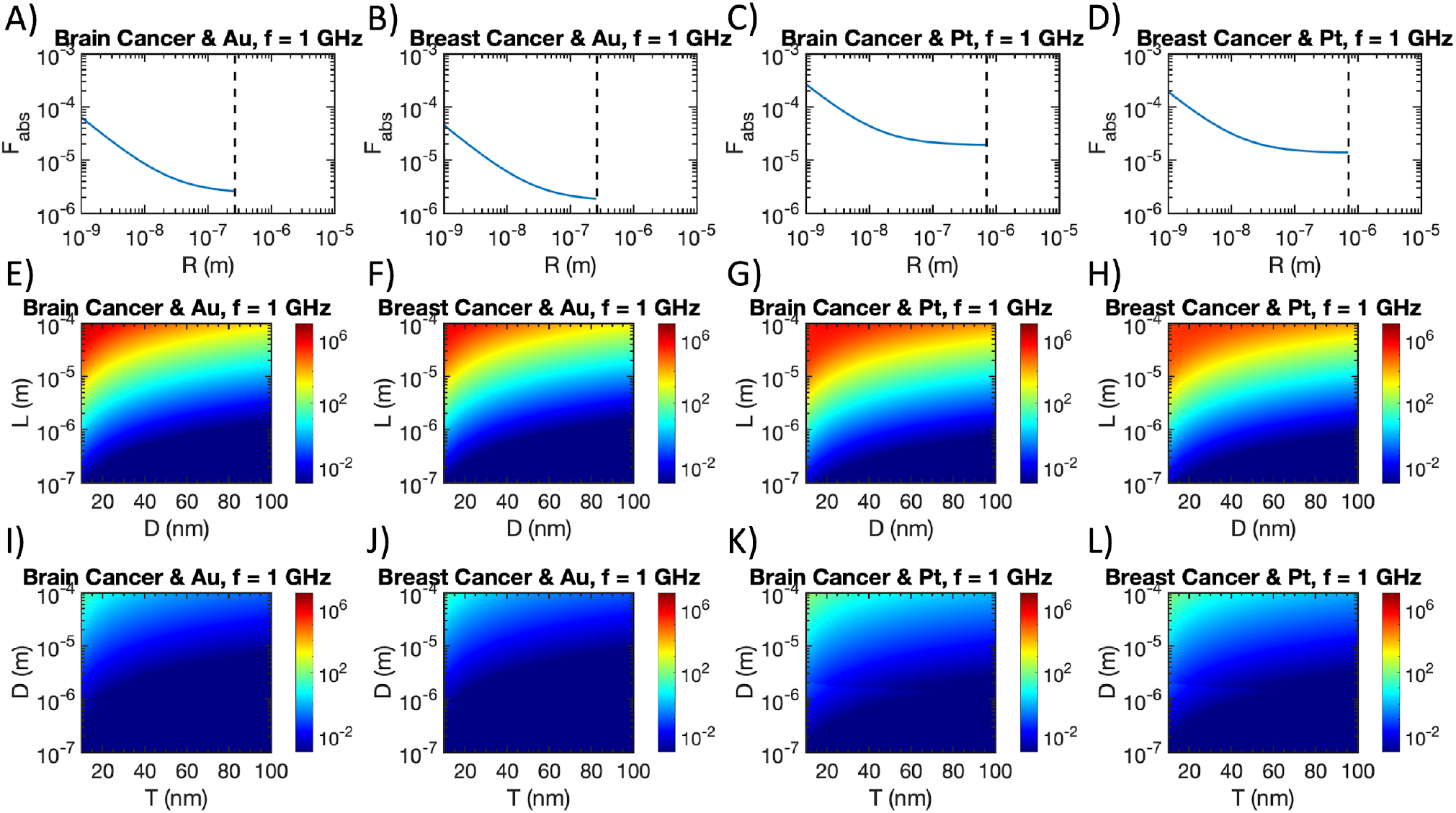
Relative absorption ratio for spherical nanoparticles (5A-5D), prolate spheroids (5E-5H), and oblate spheroids (5I-5L) in breast and brain cancerous tissues at 1 GHz. Dielectric properties of cancerous tissues are taken from Ref. [45] for breast cancer and Ref. [44] for brain cancer. Dashed lines in Figures 5A-5D represent the boundary of the electrostatic approximation.

## Discussion

The results above demonstrate that metallic nanoparticles can indeed be useful transducers of RF or microwave radiation into heat for hyperthermia treatments, but the geometry and aspect ratio of the nanoparticles are crucial to determining the degree of heating. While spherical nanoparticles are commonly synthesized in laboratories and have been extensively employed in biological applications, the absorption of spherical particles is too low when compared to a background of biological tissue. This result agrees with the theoretical and experimental investigations ruling out Joule heating of spherical gold nanoparticles in RF [19–21].

Prolate spheroids (approximating nanowires) offer powerful differential heating capable of minimizing off-target heating. Nanowires of various metals are also commonly synthesized today, with adjustable diameters and lengths, making them suitable for affordable and tunable hyperthermia treatments [55]. Based on the results of Figures 3B-3H, it is crucial to employ high aspect ratio nanowires in order to maximize *F*_*abs*_. Similar results have been reported experimentally, where silver nanowires of higher aspect ratio demonstrated more powerful heating than low aspect ratio nanowires at 13.56 MHz [29,30]. While the length of the longest prolate spheroids considered in Figure 3 may be not be suitable for all *in vivo* applications (e.g. intravenous delivery), nanowires even 1 µm in length offer a factor of 10 increase in *C*_*abs*_ over the background tissue, which could still be useful for concentrating heating in hyperthermia treatments.

Oblate spheroids (approximating nanodiscs) offered only slight differential heating in tissue, with a maximum *F*_*abs*_ of ∼10^2^. While this degree of differential heating may be suitable for some hyperthermia applications, nanodiscs are also less commonly fabricated than nanowires, and thus the practicality of this geometry may be limited when compared to nanowire-based hyperthermia treatments.

To provide practical guidance for hyperthermia researchers and clinicians, we calculate *F*_*abs*_ for various reported or commercially available nanostructures in Table 1. Specifically, we calculate *F*_*abs*_ for these nanostructures in breast and brain tissue at 1 GHz. We find that none of the spherical nanoparticles nor nanodiscs produce differential heating. Commercially available nanowires show minimal promise for differential heating under these conditions (max *F*_*abs*_∼ 10), while lab-grown nanowires reported in the literature produce *F*_*abs*_ between 10^4^ and 10^6^. This survey of realistic nanostructures demonstrates the importance of applying high-aspect-ratio nanowires for hyperthermia treatments. While Figure 4 suggests some promise of differential heating for high-aspect-ratio nanodiscs, sufficiently thin and broad nanodiscs have not yet been demonstrated in the literature, although they may be achievable with lithography and physical vapor deposition. We further guide hyperthermia researchers to the most promising regions of our parameter space in Table 2, with high-aspect-ratio prolate spheroids of Cu in breast tissue around 1 GHz offering the largest *F*_*abs*_ of the parameter space investigated in this manuscript. Our theoretical prediction (Table 2) agrees with existing experimental observations (Table 3): the lack of differential heating of spherical nanoparticles in microwave and RF fields has been confirmed in multiple papers [19,20], whereas Ag nanowires demonstrate significant differential heating in a 13.56-MHz electromagnetic field only recently [29]. Our theoretical results in Table 2 suggest a considerable room for improvement by applying 1 GHz microwaves to Cu nanowires with a diameter of 10 nm and a length of 100 μm to produce the greatest differential heating.

**Table 1:** Relative absorption ratio of various commercial or reported nanostructures.

| Material and geometry | Dimensions | Reference or vendor | $F_{abs}$ in brain tissue at 1 GHz | $F_{abs}$ in breast tissue at 1 GHz |
| --- | --- | --- | --- | --- |
| Ag nanosphere | $D = 20$ nm | Sigma Aldrich | $6.3 \times 10^{-6}$ | $1.3 \times 10^{-6}$ |
| Au nanosphere | $D = 100$ nm | Nanopartz | $3.0 \times 10^{-6}$ | $5.9 \times 10^{-7}$ |
| Ag nanowire | $D = 40$ nm, $L = 110$ μm | [62] | $1.5 \times 10^5$ | $3.3 \times 10^4$ |
| Cu nanowire | $D = 35$ nm, $L = 200$ μm | [57] | $1.6 \times 10^6$ | $7.8 \times 10^5$ |
| Au nanowire | $D = 75$ nm, $L = 10$ μm | Nanopartz | 2.0 | 0.41 |
| Au nanowire | $D = 30$ nm, $L = 6$ μm | Sigma Aldrich | 15 | 3.0 |
| Ag nanodisc | $T = 20$ nm, | [63] | $3.6 \times 10^{-4}$ | $7.1 \times 10^{-5}$ |
| | $D = 530 \text{ nm}$ | | | |
| Pt nanodisc | $T = 20 \text{ nm},$<br>$D = 530 \text{ nm}$ | [63] | $2.4 \times 10^{-3}$ | $4.8 \times 10^{-4}$ |
| Au nanodisc | $T = 20 \text{ nm},$<br>$D = 530 \text{ nm}$ | [63] | $4.2 \times 10^{-4}$ | $8.2 \times 10^{-5}$ |
| Au nanodisc | $T = 35 \text{ nm},$<br>$D = 800 \text{ nm}$ | [64] | $1.2 \times 10^{-4}$ | $2.4 \times 10^{-5}$ |

**Table 2:** Optimal nanostructures to maximize relative absorption ratio.

| Geometry | Optimal conditions |
| --- | --- |
| Spherical particles | Differential heating does not occur under any conditions. |
| Prolate spheroids | The greatest differential heating occurs around 1 GHz for prolate spheroids made of Cu with $D = 10 - 20 \text{ nm}$ and $L = 100 \text{ }\mu\text{m}$ in breast tissue. Note that the optimal size parameters are limited by the choice of the parameter space in Figure 3. However, a general trend to maximize $F_{abs}$ holds: a smaller diameter $D$ and a greater length $L$ result in a higher $F_{abs}$ for prolate spheroids composed of any metal in any tissue. In the high aspect-ratio limit, the optimal frequency $f$ decreases with an increasing length $L$. $F_{abs}$ is slightly dependent on metal and tissue types, with Cu in breast tissue demonstrating the greatest $F_{abs}$ in the parameter space of interest. |
| Oblate spheroids | There is minor potential for differential heating between 1-10 GHz for oblate spheroids with $T = 10 \text{ nm}$ and $D > 10 \text{ }\mu\text{m}$. Such nanodiscs have not yet been demonstrated but may be attainable with lithography and physical vapor deposition. |

**Table 3:**
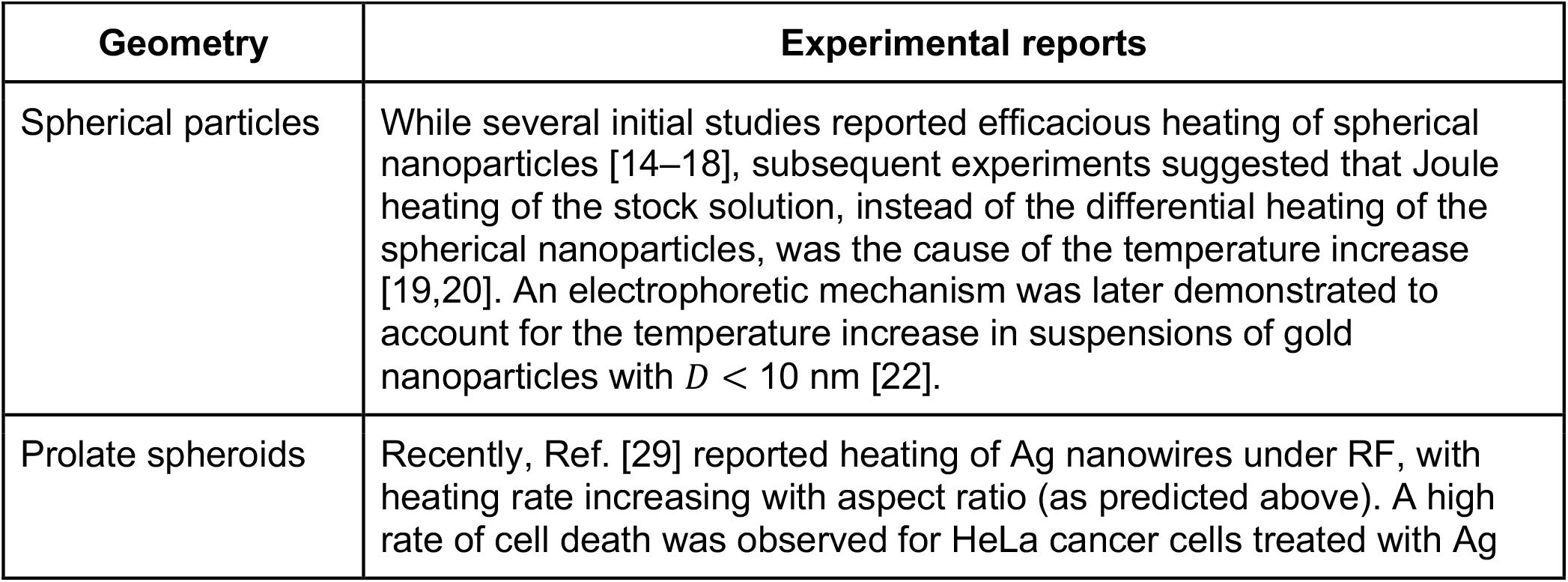
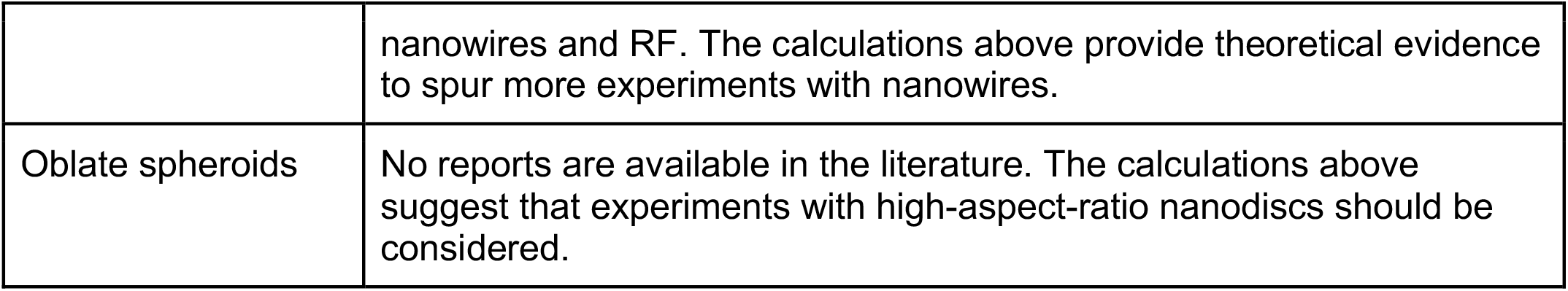
Previous experimental reports of hyperthermia with various nanostructures.

| Geometry | Experimental reports |
| --- | --- |
| Spherical particles | While several initial studies reported efficacious heating of spherical nanoparticles [14–18], subsequent experiments suggested that Joule heating of the stock solution, instead of the differential heating of the spherical nanoparticles, was the cause of the temperature increase [19,20]. An electrophoretic mechanism was later demonstrated to account for the temperature increase in suspensions of gold nanoparticles with $D < 10 \text{ nm}$ [22]. |
| Prolate spheroids | Recently, Ref. [29] reported heating of Ag nanowires under RF, with heating rate increasing with aspect ratio (as predicted above). A high rate of cell death was observed for HeLa cancer cells treated with Ag |
|  | nanowires and RF. The calculations above provide theoretical evidence to spur more experiments with nanowires. |
| Oblate spheroids | No reports are available in the literature. The calculations above suggest that experiments with high-aspect-ratio nanodiscs should be considered. |

As an example of the temperature increase that may be expected from realistic nanostructures considered in this manuscript, we consider a 2 µL bolus injection (typical viral injection volume [56]) of the Cu nanowires reported in Ref. [57] at a volume fraction of 10^−3^. With *F*_*abs*_ = 1.6 × 10^6^ for these nanowires (Table 1), we expect an effective *F*_*abs*_ = 1.6 × 10^3^ for the bolus injection. We solve the Pennes bioheat equation (see **Supplementary Information** for methods) in brain tissue over 10 s of heating with a 1 GHz field applied at 300 V/m. We selected this field strength because it heats non-injected brain tissue by only 0.1°C over 10 s, avoiding damage to surrounding healthy tissue. Meanwhile, the center of the Cu nanowire bolus injection heats by approximately 26°C over 10 s, providing more than enough heat to induce local tumor cell death [3].

While much of the literature thus far has focused on spherical gold nanoparticles, the results presented above illuminate the opportunities afforded by alternative geometries and materials. By working within the bounds of the electrostatic approximation, analytic expressions can be used to calculate the relative absorption ratio across a broad parameter space, minimizing computational cost and enabling straightforward optimization of hyperthermia techniques. Although Joule heating has been repeatedly discounted in the case of spherical nanoparticles, this analysis suggests that current flow within non-spherical particles can indeed produce meaningful differential heating. Compared to previous reports that present analytical and numerical solutions of anisotropic nanoparticles to afford maximum differential heating in a lossy medium [27,32], this work is the first to demonstrate the feasibility of RF/microwave-based remote hyperthermia treatments *in vivo* with realistic metal nanostructures. Furthermore, our results provide a practical guide for choosing the material, size, and geometry of nanoparticles to afford desired local hyperthermia therapy in biological tissues *in vivo* under RF/microwave fields of various frequencies.

## Conclusion

We have analytically examined the relative absorption ratio of metallic nanostructures in biological tissue based on an array of theoretical frames including the classical electromagnetic theory, the Drude model, and the electrostatic approximation. We have confirmed that spherical nanoparticles do not offer differential heating in tissue, while higher-aspect-ratio geometries such as nanowires (and to a lesser extent, nanodiscs) offer strong absorption of RF and microwave radiation. Slight differences in the maximum absorption occur depending on the metal and tissue type considered, but the above calculations can be easily performed in the clinic for optimization as needed. We believe these results are immediately relevant to hyperthermia treatments, and subsequent work will examine the attainable increase in tissue temperature both around individual nanoparticles and in the bulk volume around a nanoparticle injection, based on the relative absorption ratios calculated in this manuscript.

## Acknowledgements

G.H. acknowledges the support by a National Institutes of Health (NIH) Pathway to Independence Award (National Institute on Aging 5R00AG056636-04), a National Science Foundation (NSF) CAREER Award (2045120), a gift from the Spinal Muscular Atrophy (SMA) Foundation, and seed grants from the Wu Tsai Neurosciences Institute and the Bio-X Initiative of Stanford University.

## Supplementary Information

### I. Confirming the validity of the Drude model at low frequencies

Consider the Drude model for free electrons given in Equation (1). We begin by extracting the real and imaginary parts of the dielectric function:

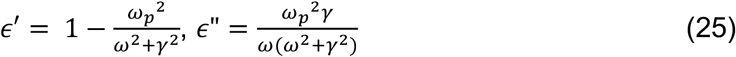

Taking the low frequency limit *ω* ≪ *γ*:

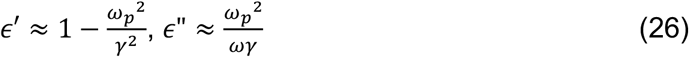

In a free electron model (assuming minimal contribution from bound electrons), ℜ(*σ*) = *ϵ*_0_*ϵ*″*ω* [31], so at low frequencies:

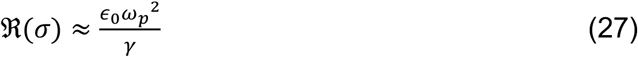

As *ω* → 0 for an applied electric field, the induced current is in phase with the applied field [58], so 𝔍 (*σ*) → 0 and 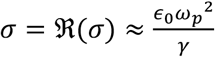. Rewriting *ω*_*p*_ using its constituent factors (defined in Equation 2) and *γ* as *τ*^−1^ where *τ* is the characteristic scattering time, we obtain:

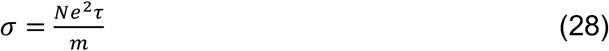

which matches the definition of the DC conductivity of metals *σ*_*DC*_ [58]. Therefore, the Drude model is valid at the low frequency limit and can thus be applied to RF frequencies. The validity of the Drude model as a good fit to the dielectric function of metals at low frequencies has also been experimentally proved [34].

### II. Choice of input parameters for the Drude model

In the calculations above, we substituted values of the input parameters *ω*_*p*_ and *γ*_*bulk*_ reported for Au, Ag, Pt, and Cu by reference [59]. We have reproduced these values in Table S1 below for convenience. However, values of *ω*_*p*_ and *γ*_*bulk*_ reported in the literature vary, and the choice of these inputs to the Drude model ultimately affects < *C*_*abs*_ >. For example, Table 1 in reference [60] gathers unique sets of *ω*_*p*_ and *γ*_*bulk*_ for gold reported by seven different studies.

To confirm the conclusions above, in Figure S1 we reproduce several key subplots for gold nanostructures from Figures 2-4, but we display the plots corresponding to the lowest and highest differential heating in the parameter space of interest. For example, Figures S1A and S1D cover the same parameter space as Figure 2B for spherical gold nanoparticles in the pancreas. We calculated seven maps of this parameter space, with each map using a unique set of *ω*_*p*_ and *γ*_*bulk*_ reported in reference [60]. We then determined the lowest and highest cases of differential heating by ranking all seven maps by the maximum *F*_*abs*_ occurring within each graph. The lowest-ranking map (minimum differential heating) is shown in Figure S1A, and the highest-ranking map (maximum differential heating) is shown in Figure S1D. These maps put bounds on the expected absorption properties for spherical gold nanoparticles in the pancreas, as an illustrative case for how variations in input parameters may affect the conclusions of this manuscript. We repeated this process for gold prolate spheroids in breast tissue (covering the parameter space of Figure 3I) in Figures S1B and S1E and gold oblate spheroids in breast tissue (covering the parameter space of Figure 4I) in Figures S1C and S1F. We observe minimal differences between the plots when comparing the minimum and maximum differential heating for spheres, prolate spheroids, and oblate spheroids, suggesting that variation of reported *ω*_*p*_ and *γ*_*bulk*_ in the literature does not significantly affect our conclusions above.

### III. Differential heating calculations

We solved the three-dimensional Pennes bioheat equation in homogenous brain tissue using the bioheatExact function in the k-Wave MATLAB toolbox [61]. Thermal properties of the brain and blood were taken from Ref. [6]. We used a 1 cm^3^ cubic domain (designed to approximate the brain of a mouse) with a uniform initial temperature distribution of 37°C, and a wave with frequency 1 GHz and strength 300 V/m that minimized heating of surrounding tissues.

**Table S1:** Input parameters for the Drude model, from reference [59].

| Metal | $\omega_p$ (PHz) | $\gamma_{bulk}$ (THz) |
| --- | --- | --- |
| Au | 2.18 | 6.45 |
| Ag | 2.18 | 4.35 |
| Pt | 1.24 | 16.7 |
| Cu | 1.79 | 2.19 |

**Figure S1:**
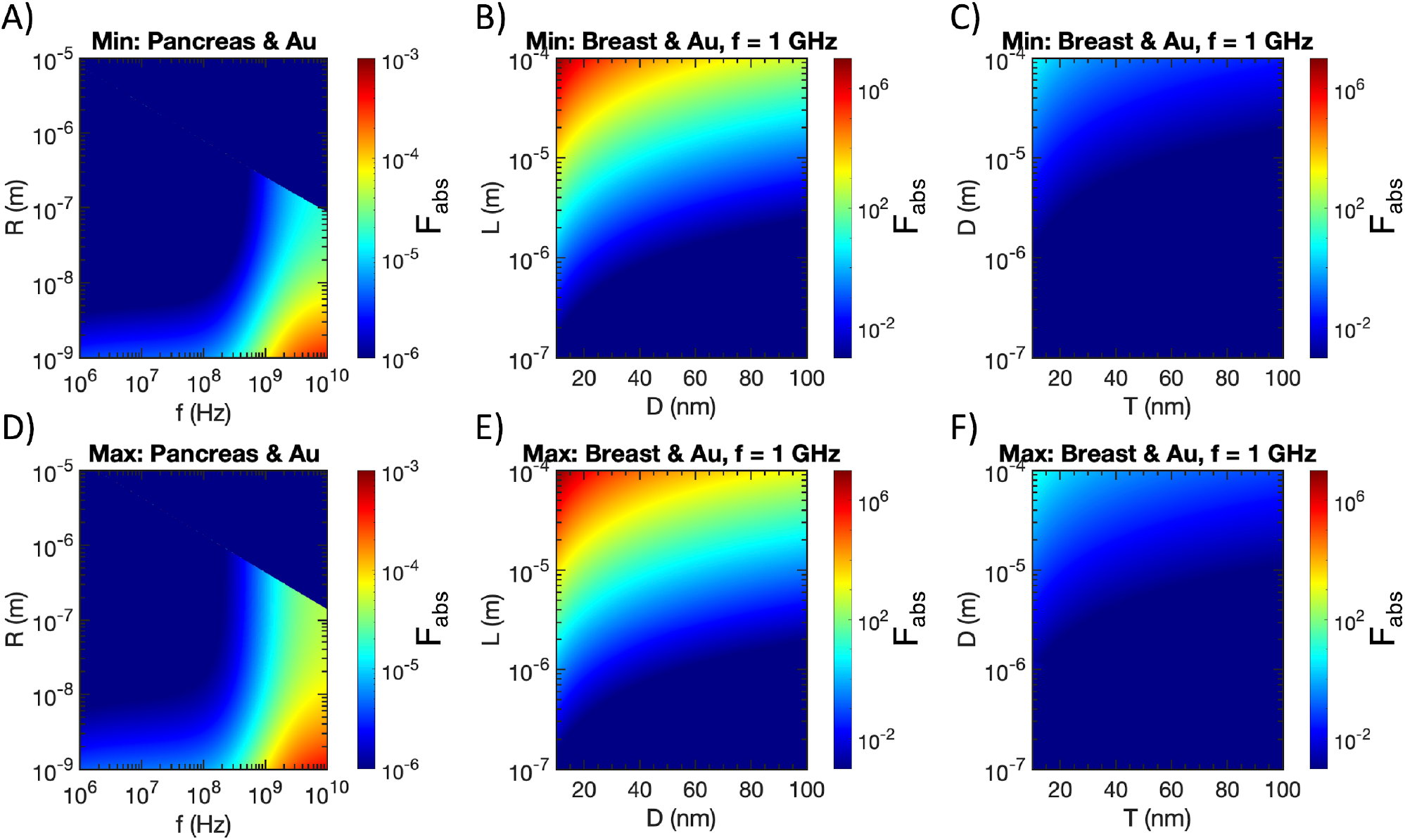
Lowest (minimum) case of differential heating for spherical gold particles in the pancreas (A), prolate gold spheroids in breast tissue (B), and oblate gold spheroids in breast tissue (C), obtained by calculations with the 7 pairs of *ω*_*p*_ and *γ*_*bulk*_ reported in Table 1 of reference [60]. Highest (maximum) case of differential heating for spherical gold particles in the pancreas (D), prolate gold spheroids in breast tissue (E), and oblate gold spheroids in breast tissue (F), obtained by calculations with the 7 pairs of *ω*_*p*_ and *γ*_*bulk*_ reported in Table 1 of reference [60]. Semi-axes that violate the electrostatic approximation are excluded from all plots.

